# Pseudouridine prevalence in Kaposi’s sarcoma-associated herpesvirus transcriptome reveals an essential mechanism for viral replication

**DOI:** 10.1101/2023.01.31.526461

**Authors:** Timothy J. Mottram, Katherine L. Harper, Elton J. R. Vasconcelos, Chinedu A. Anene, Adrian Whitehouse

## Abstract

Pseudouridylation is a prevalent RNA modification shown to occur in tRNAs, rRNAs, snoRNAs and most recently mRNAs and lncRNAs. Emerging evidence suggests that this dynamic RNA modification is implicated in altering gene expression by regulating RNA stability, modulating translation elongation and modifying amino acid substitution rates. However, the role of pseudouridylation in infection is poorly understood. Here we demonstrate that Kaposi’s sarcoma-associated herpesvirus (KSHV) manipulates the pseudouridylation pathway to enhance replication. We show the pseudouridine synthases (PUS), PUS1 and PUS7 are essential for efficient KSHV lytic replication, supported by the redistribution of both PUS1 and PUS7 to viral replication and transcription complexes. We present a comprehensive analysis of KSHV RNA pseudouridylation, revealing hundreds of modified RNAs at single-nucleotide resolution. Notably, we further demonstrate that pseudouridylation of the KSHV-encoded polyadenylated nuclear RNA (PAN) plays a significant role in the stability of PAN RNA and in the association of the KSHV ORF57 protein. Our findings reveal a novel and essential role of pseudouridine modification in the KSHV replication cycle.

## Introduction

Post-transcriptional chemical modifications of RNAs are widely abundant across all forms of RNA, affecting up to 25% of all nucleotides present. There are over 170 modifications currently identified, exhibiting a plethora of functions including RNA stabilisation, localisation and the facilitation of intermolecular interactions. Resurging interest in RNA modifications has been driven by advancements in transcriptome-wide RNA modification mapping and the identification of RNA modifications in all RNA species, including mRNA and ncRNAs. Pseudouridine (Ψ) is the most abundant single nucleotide modification found in all functional RNA species ^1, 2^. Ψ is catalysed by two groups of enzymes, RNA dependent (H/ACA Box snoRNA-guided) such as Dyskerin ^3, 4, 5^, or RNA independent (direct) known as pseudouridine synthases (PUSs) ^6, 7^. PUS enzymes function to break the carbon-nitrogen bond found in uridine, then subsequently reform a carbon-carbon bond through the C5 position of the cleaved uridine to the ribose sugar. This function can be site dependent, driven by specific motif binding or secondary RNA structure ^6^. The function of Ψ within RNA species such as tRNA, snoRNA and rRNAs are well characterised, however functions of Ψ within mRNA and ncRNA is largely unknown. Recent transcriptome-wide studies have revealed Ψ can be dynamically modified in response to cellular stress, which may unveil an analogous function to other modifications such as m^6^A. Ψ has confirmed functions in RNA folding, protein binding, protein translation, RNA-RNA interactions and RNA stability ^8, 9, 10, 11^, however evidence suggests this is a transcript specific, rather than global effect, as often when Ψ is removed, stability of the RNA molecule remains intact. Changes to Ψ-modified mRNA status often occur when under cell stress and confer an enhancement to cell survivability ^12, 13^. The identification of how Ψ sites function in host or pathogen mRNAs and ncRNAs during infection is currently understudied.

RNA modifications such as m^6^A are found in a wide range of viruses. The ability of such modifications to regulate gene expression offers unique possibilities for viruses to modulate viral and host genes, but also for the host to regulate a response to infection ^14^. For example, m^6^A has been identified on transcripts encoded by a wide range of viruses and studies to investigate m^6^A function in virus life cycles have highlighted distinct roles indicating widespread regulatory control over viral life cycles ^15^. Additionally, m^5^C, a modification prevalent across many virus genomes, may play a role in modulating viral gene expression. For instance, m^5^C knockdown in HIV-1 resulted in dysregulation of alternative splicing within viral RNAs ^16^. Most recently, Ψ was identified within Epstein-Barr virus (EBV)-encoded non-coding RNA EBER2, proving essential for the stability of the RNA and required for efficient lytic viral replication ^17^. However, little is known of Ψ influence on regulatory mechanisms controlling virus replication and a lack of global transcriptome-wide analysis has yet to unveil the prevalence of Ψ within viral genomes.

Kaposi’s sarcoma-associated herpesvirus (KSHV) is a large double stranded DNA virus associated with Kaposi’s sarcoma and two lymphoproliferative disorders: primary effusion lymphoma and multicentric Castleman’s disease. Like all herpesviruses, KSHV has a biphasic life cycle comprising of latent persistence and lytic replication cycles. KSHV establishes latency in B cells and in the tumor setting, where viral gene expression is limited to a small subset of viral genes allowing the viral genome to persist as a non-integrated episome. Upon reactivation through certain stimuli such as cell stress; KSHV enters the lytic replication phase, leading to the orchestrated temporal expression of over 80 viral proteins necessary for the production of infectious virions ^18^. Notably, both the latent and lytic replication phases are essential for KSHV-mediated tumorigenicity ^19^. Interestingly several recent studies have shown that m^6^A is highly prevalent throughout the KSHV transcriptome and enhances the stability of the essential latent-lytic switch transcriptional protein RTA transcript ^20, 21^. This highlights the importance of RNA modifications in regulating viral gene expression.

Herein we use a quantitative proteomic approach to identify changes in PUS1 and PUS7 enzymes interactome during KSHV lytic replication. Furthermore, CRISPR-Cas9 knockout analysis of PUS1 and PUS7 confirmed they are essential for KSHV lytic replication. Subsequent RBS-Seq analysis demonstrates that the KSHV transcriptome is functionally pseudouridylated, with over 200 candidate Ψ sites. Notably, specific Ψ sites play an important functional role in the polyadenylated nuclear RNA (PAN), by allowing ORF57 to enhance PAN expression through changes to RNA stability. These results describe a novel mechanism for KSHV to utilise host cell post-transcriptional RNA modification machinery to modify KSHV RNA transcripts enabling efficient viral replication.

## Results

### PUS enzymes are essential for KSHV lytic replication

To determine whether KSHV manipulated the host cell pseudouridylation machinery we first sought to identify any changes in localisation of the PUS enzymes, specifically PUS1 and PUS7 during KSHV reactivation. TREx BCBL1-Rta cells, a KSHV-latently infected B-lymphocyte cell line containing a Myc-tagged version of the viral RTA under the control of a doxycycline-inducible promoter, remained latent or were reactivated for 24 hours prior to immunostaining with antibodies against KSHV early proteins and PUS1 or PUS7 proteins. PUS1 showed a diffuse staining throughout the nucleus and cytoplasm in latently infected cells. During lytic replication, nuclear PUS1 was redistributed to the replication and transcription compartments (RTCs), a virally-induced intra-nuclear structure where viral transcription, viral DNA replication and capsid assembly occur (Fig. 1a). In contrast, PUS7 was redistributed from a diffuse nuclear localisation exclusively into KSHV RTCs (Fig. 1b).

**Figure 1.**
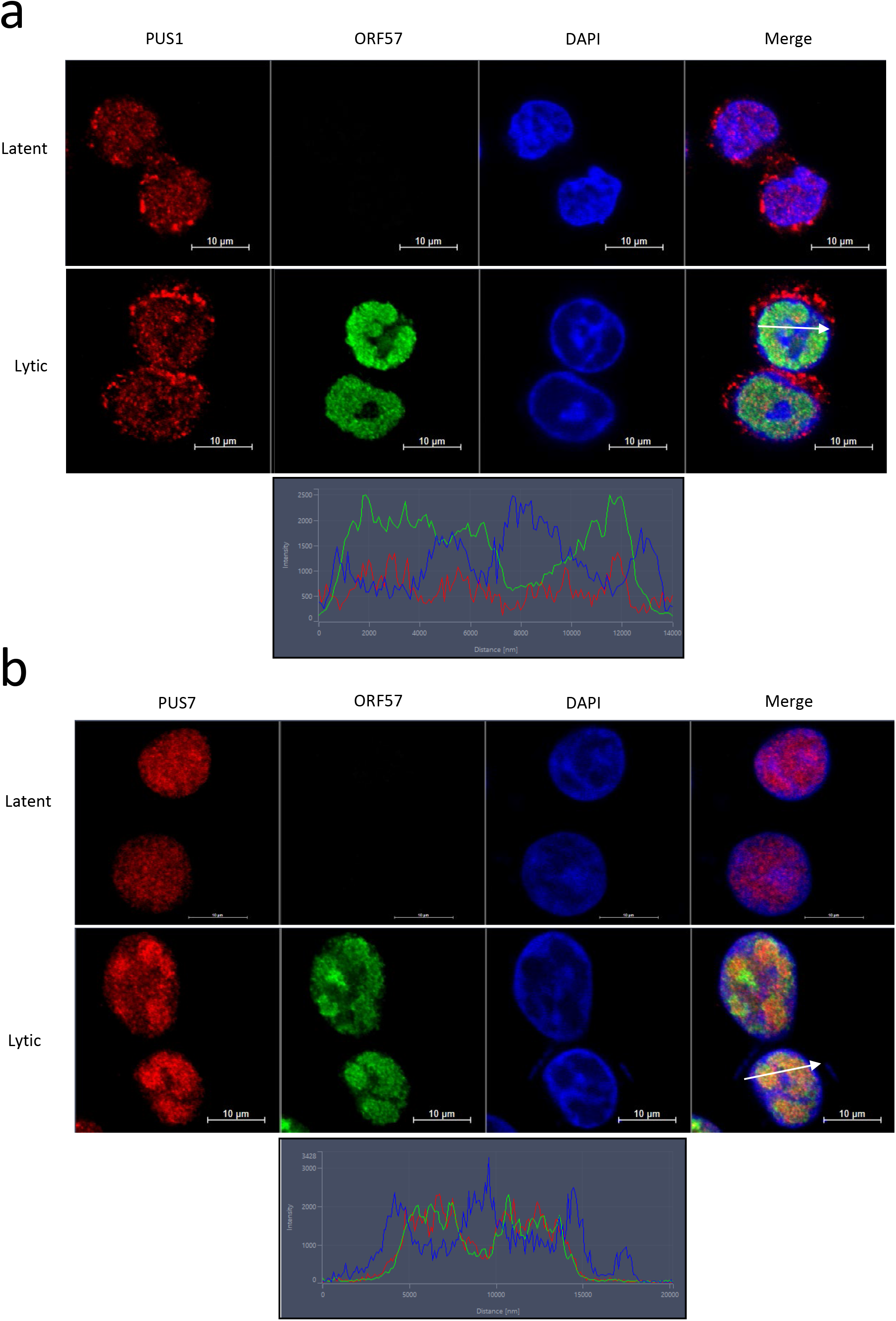
PUS1 and PUS7 are essential for KSHV viral replication and PUS7 shows distinct relocalisation upon viral reactivation. TREx BCBL1-Rta cells were reactivated using doxycycline and fixed at 24 hours post activation. Untreated TREx BCBL1-Rta cells were used as a control. Cells were stained using anti-PUS1 (a) or anti-PUS7 (b) to stain for PUS localisation and anti-ORF57 antibodies to identify replication centres. Cross section fluorescent intensity graph of representative cell identified by a white arrow. Representative image of n = 3 shown.

To determine whether PUS enzymes were essential for KSHV lytic replication, PUS1 (Fig. 2a) and PUS7 (Fig. 2b) were successfully knocked out in TREx BCBL1-Rta cells utilising the lentiCRISPRv2 CRISPR-cas9 system. To ensure the CRISPR cas9 single cell cloning did not result in large variations in viral episome count, assessment of latent viral DNA was performed (Supplementary Fig. 1), which confirmed single cell clones had no significant changes to KSHV episome load. Reactivation of PUS1ko and PUS7ko showed a reduction in early KSHV ORF57 protein levels, however there was a complete abolition of late ORF65 protein production (Supplementary Fig. 2a-b), suggesting pseudouridylation may be important in the later stages of the KSHV lytic temporal cascade. Furthermore, viral RNA expression of immediate-early gene PAN (Fig. 2c), early gene ORF57 (Fig. 2d) and late gene ORF65 (Fig. 2e) ^18^ was impaired, with PAN and ORF57 showing a significant ~30% reduction and ORF65, a further ~80%, indicating that the effect seen is at a transcriptional level. To confirm that knockout of PUS1 and PUS7 affected infectious virion production, supernatants of reactivated PUS1ko or PUS7ko TREx BCBL1-Rta cells were used to re-infect naïve HEK-293T cells and KSHV ORF57 expression was determined by qPCR (Fig. 2 f-g). Cells reinfected with supernatant from both PUS1ko and PUS7ko showed a significant reduction (>90%) in infectious virion production. Together this suggests that KSHV may utilise PUS enzymes, redistributing both PUS1 and PUS7 into RTCs, to enhance later stages of viral lytic replication.

**Figure 2.**
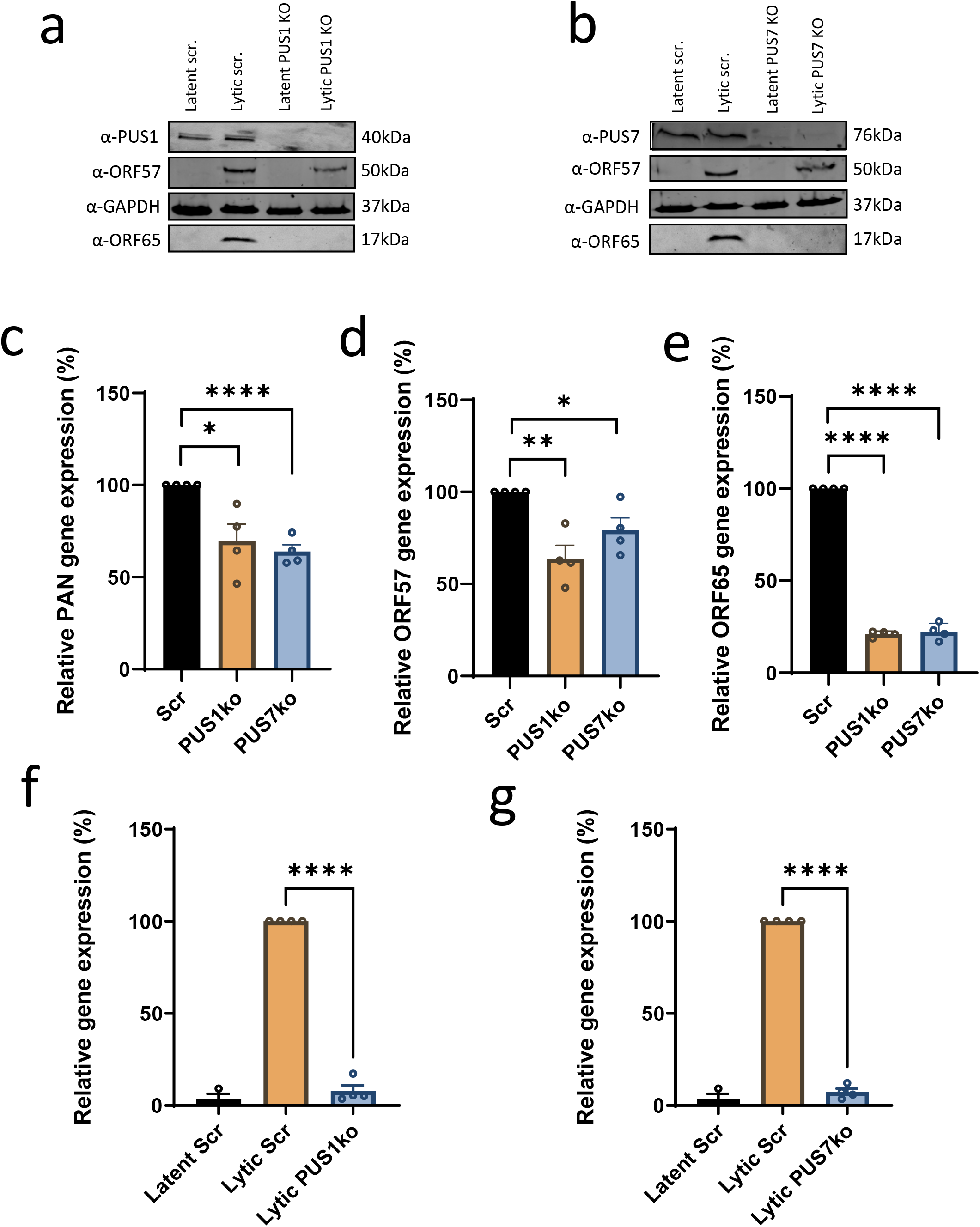
Early and late viral RNA and protein expression of reactivated KSHV are disrupted in PUS1ko or PUS7ko cells. Early and late viral protein expression of reactivated KSHV in PUS1ko (a) or PUS7ko cells (b). Expression of viral early protein ORF57, late protein ORF65 and housekeeping protein GAPDH in TREx BCBL1-Rta cells as determined by Western blot using anti-PUS1 or anti-PUS7, ORF57, ORF65 and GAPDH specific antibodies. Representative image of three biological repeats shown. Gene expression of immediate-early gene non-coding RNA PAN (C), early gene ORF57 (D) and late gene ORF65 (E) in TREx BCBL1-Rta PUS1ko or PUS7ko cells after 24 hours lytic reactivation. Viral RNA levels determined via qRT-PCR and normalised to GAPDH. Values were normalised to scrambled control KSHV infection Error bars represent SD, n = 4 for all experiments, p ≤ 0.05 *, p ≤ 0.01 **, p ≤ 0.0001 **** using a two tailed Students unpaired t test. Reduced production of infective of KSHV virions in PUS1ko (F) or PUS7ko (G) cells. Successful infection, and replication of KSHV virions was determined by reinfection of naïve HEK-293Ts. After 72 hours of doxycycline induction in TREx BCBL1-Rta cells, supernatant was used to infect HEK-293Ts, which were subsequently harvested at 48 hours post infection. Viral mRNA levels were determined via qRT-PCR of the ORF57 gene and normalised to GAPDH. Values were normalised to scrambled control KSHV infection. Error bars represent SD, n = ≥ 3 for all experiments, p ≤ 0.0001 **** using a two tailed Students unpaired t-test.

### Pus enzyme interactomes are altered during KSHV lytic replication

To determine whether the KSHV-mediated redistribution of PUS1 and PUS7 into RTCs affects PUS enzyme protein-protein interactions, affinity pulldowns were performed using anti-PUS1, anti-PUS7 or control anti-IgG antibodies in latent or reactivated KSHV-infected cells, prior to analysis by TMT-labelled quantitative mass spectrometry. Negative control (anti-IgG pulldown) protein abundance was subtracted from total abundance for each protein followed by a fold change comparison between KSHV latent and lytic samples. Samples with a total protein abundance below 100 for any condition were discarded. The change in interaction landscape of both PUS1 and PUS7 during lytic replication is consistent with the observed relocalisation as shown by immunofluorescence. PUS1 showed an upregulation in interactions with proteins involved in biological processes such as cell division, molecular chaperones and translational factors (Fig. 3a) including important cellular proteins EEF1A1, HSP60 and HSP90, suggesting the redistribution of PUS1 to RTCs facilitates new protein-protein interactions. Notably, STRING analysis suggested that relocalisation of PUS7 resulted in a significant down regulation in interactions with cellular proteins. These include ribonucleoproteins such as hnRNP C1/2, hnRNP A2/B1 and hnRNP U along with nuclear, translational and histone factors, such as H2B, H1.4 and H4. All interactions lost are known PUS7 interaction groups under stable cellular conditions (Fig. 3b) ^22^. Differentially expressed PUS1 and PUS7 interaction correlation and hierarchical clustering analyses with whole cell TREx BCBL1-Rta SILAC proteomics that allowed assessment of global protein expression levels during lytic reactivation (Fig. 3c) revealed limited correlation (rho <0.2), confirming PUS1 and PUS7 increased or decreased interactions are not due to decreased overall protein expression by viral host cell shutoff.

**Figure 3.**
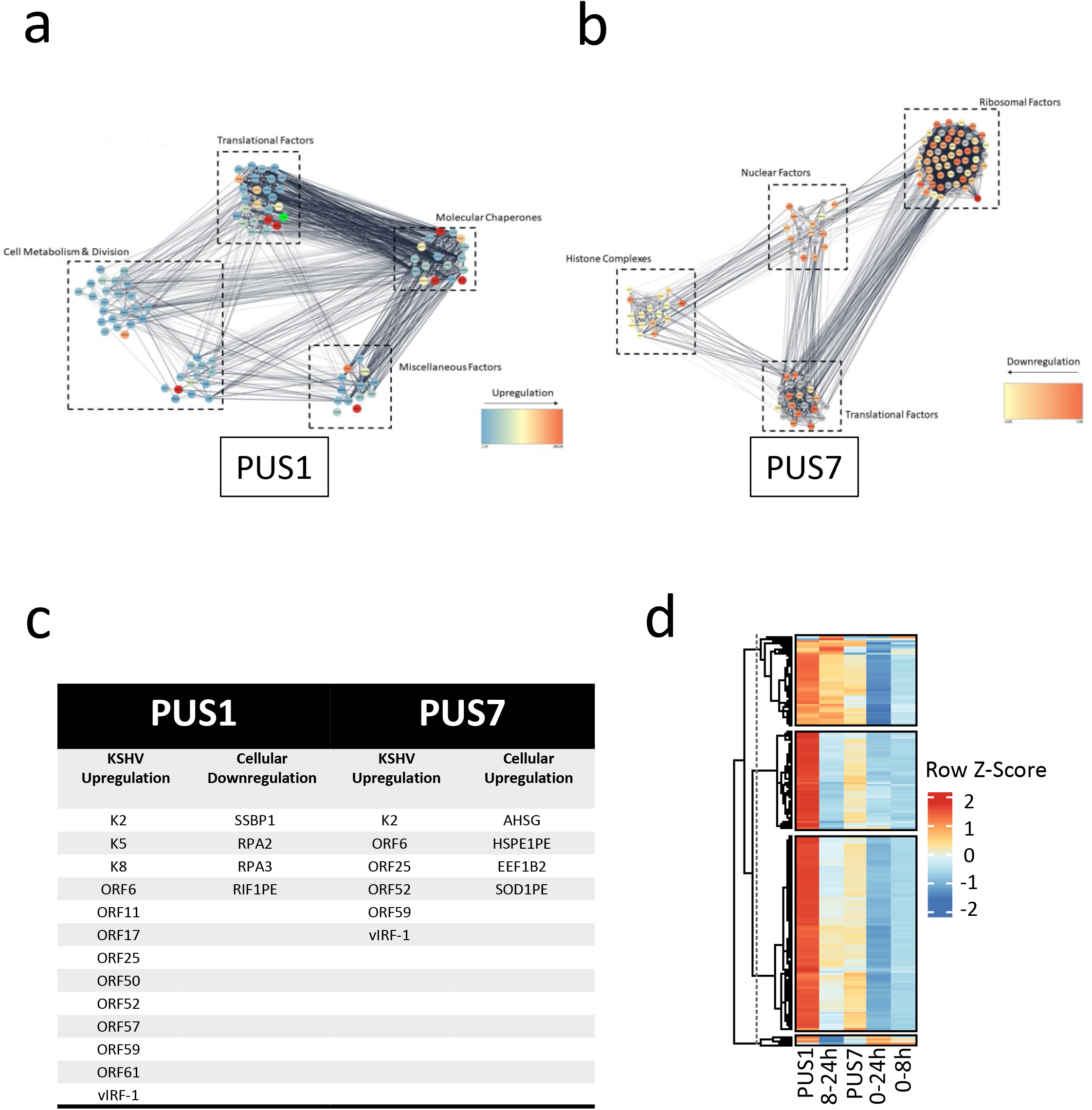
PUS1 and PUS7 enzymes show distinct changes in protein interaction partners upon KSHV lytic reactivation. STRING analysis of PUS1 upregulated (a) and PUS7 downregulated (b) interaction partners. Reactivated TREx BCBL1-RTA cells after 24 hours were harvested and an immunoprecipitation performed using anti-PUS1 or anti-PUS7 antibodies. Samples were then processed using tandem-mass tag proteomics. STRING analysis displays upregulation of cellular protein interactions with PUS1 and downregulation of interactions with PUS7. Heat map hierarchical clustering (HC) analysis between differentially expressed proteins during KSHV reactivation and PUS1/PUS7 immunoprecipitation interaction partners (d). HC was performed on both rows (proteins) and columns (samples), and the colour shading scale corresponds to Z-Scores with red colour indicating higher interaction and blue colour indicating lower interaction. Spearman correlation coefficient (rho) was also calculated on all possible pairwise comparisons of TREx BCBL1-RTA PUS1 and PUS7 immunoprecipitation data with reactivated lytic TREx BCBL1-RTA whole cell SILAC proteomics data (rho < 0.2, not shown). Schematic of PUS7 interaction hypothesis (d). PUS7s relocalisation to replication complexes brings PUS7 in close proximity to viral proteins, cellular factors and the PAN RNA scaffold, along with any transcribing viral RNAs while reducing latent cellular interactions.

TMT proteomics was assessed for differential expression commonality between PUS1 and PUS7 interaction partners (Supplementary Fig. 3). However, as expected, the divergent functions of PUS1 and PUS7 resulted in the vast majority of interactors showing no commonality. Notably however, viral proteins K2, ORF6, ORF25, ORF52, ORF59 and vIRF-1 were all identified as potential interaction partners for both PUS1 and PUS7 highlighting these interactions may have functional relevance to the pseudouridylation of viral RNA. Together, this data highlights the change in both PUS1 and PUS7 interactomes may have an impact on both cellular and viral RNA pseudouridylation status. Furthermore, we hypothesise the change in PUS1 and PUS7 localisation and resulting change in protein interactors indicates the involvement of PUS enzymes in viral RTC activity (Fig. 3d).

### Transcriptome-wide mapping of Ψ during KSHV lytic replication

To elucidate the landscape of Ψ in the KSHV transcriptome, RNA-bisulfite sequencing (RBS-Seq) was performed in latent TREx BCBL1-Rta cells and cells undergoing lytic replication for 8 h and 20 h post-induction. RBS-Seq results in a highly reproducible 1-2 nucleotide deletion signature at Ψ sites, exclusively in bisulfite (BS) treated samples. Utilising the custom bioinformatics pipeline developed by Khoddami et al ^23^, we identified 462 unique Ψ sites within the KSHV transcriptome (Fig. 4a). Of these sites, 33 were detected only during latency, associated with latently expressed transcripts, such as LANA (Fig. 4b). At 8 hours post reactivation, 160 sites were identified during the early stages of the lytic cascade. As the lytic temporal cascade progressed, 115 sites unique at 20 hours post reactivation were identified, not surprisingly 88 sites were conserved between 8 and 20 hours post reactivation, typically within lytic genes predominantly expressed throughout lytic replication (Fig. 4c). Furthermore, mapping the Ψ sites to genome features previously identified in KSHV ^18^, showed that Ψ is found predominantly within the CDS of KSHV genes with the remaining Ψ sites identified in the UTR and repeating regions (Fig. 4d). The proportional pseudouridylation of these gene features was consistent through 0, 8 and 20 hours post reactivation. By performing STREME motif analysis of 15 nt sequences flanking each Ψ site, we observed varying consensus motifs between genome features (Fig. 4e). Within the CDS, a motif of AAG was most common, which corresponds to the HRU PUS1 motif in the reverse orientation. Interestingly, a CCCMCAYCCC was the most prominently conserved motif within the intergenic region which has some similarity to human TRUB1 GUUCNANNC motif. Analysis of all Ψ site motifs showed ~49% contained either AGGAR or AAAA motifs. Furthermore, assessing the PUS7 motif UG<u>U</u>AR showed 6 sites within the CDS that are strong candidates for Ψ modification (Fig. 4f). Together this data shows that the KSHV transcriptome is heavily pseudouridylated throughout lytic reactivation.

**Figure 4.**
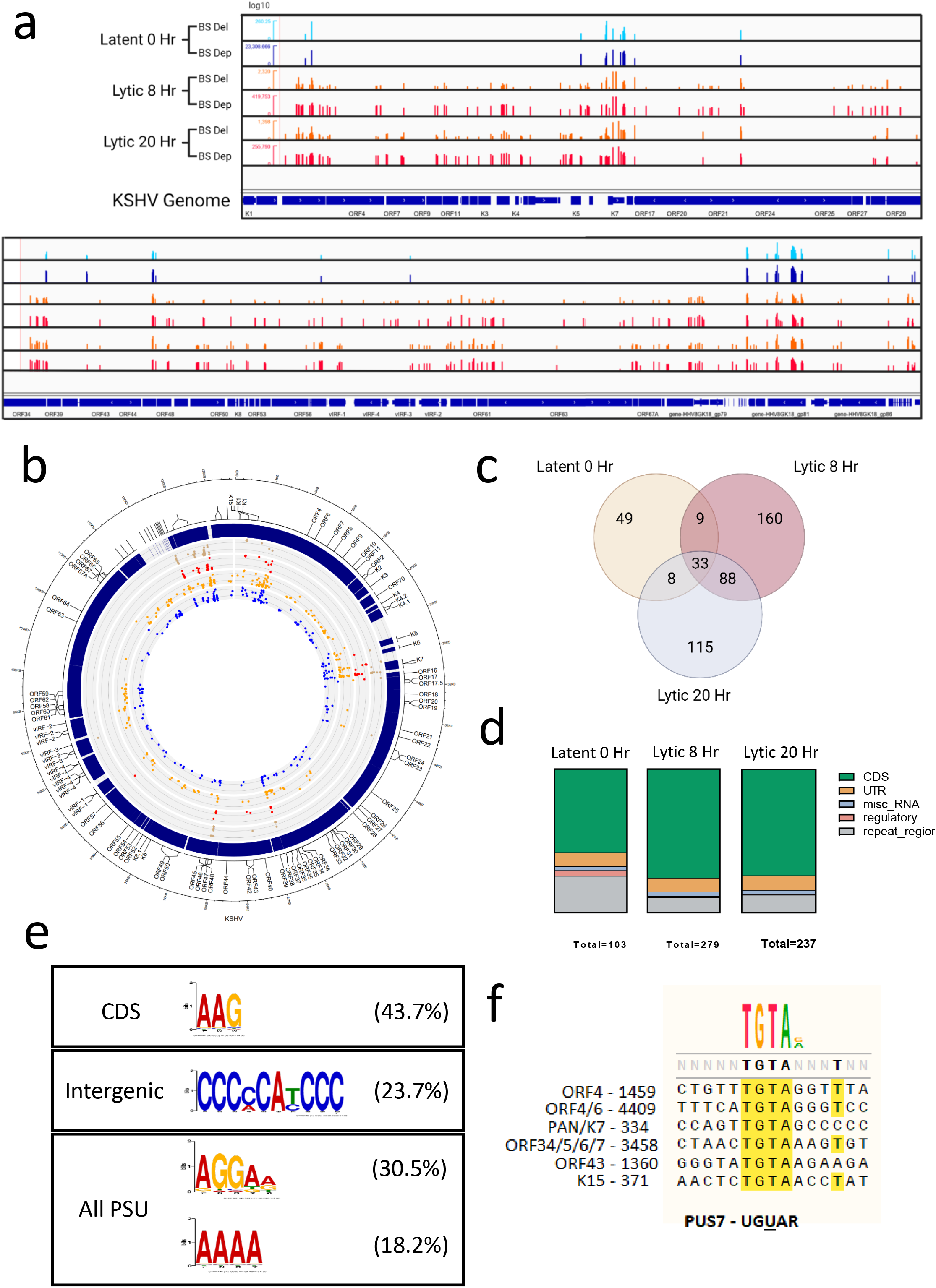
The KSHV transcriptome is heavily φ modified. RBS-seq IGV KSHV epitranscriptome map (a). Raw sequencing reads were processed as described in methods. RBS-Seq Integrative Genomics Viewer (IGV) plot in TREx BCBL1-Rta cells at 0, 8 and 20 hr post reactivation. Log10 bisulfite deletions (BS Del) and bisulfite depth (BS Dep) aligned across the KSHV epitranscriptome are shown. RBS-Seq CIRCOS analysis of individual φ sites mapped to KSHV transcriptome (b). Plot of φ sites identified by RBS-Seq analysis at 8 hours unreactivated (red), post reactivation (blue), 20 hours unreactivated (brown) and post reactivation (orange). Coverage tracks scaled to log10. Individual repeats shown. Venn diagram comparison of related φ sites during lytic reactivation (c). Comparison of overlapping φ sites at 0, 8 and 20 hr post reactivation. Distribution of φ sites across topological regions of viral RNA (d). Genome features include CDS, UTR, miscellaneous RNA, regulatory RNA and repeat regions. STREME analysis of φ motif sequences (e). Analysis of sequence motifs of within 15 nt surrounding φ site separated by genomic features (CDS and intergenic) or considering all φ sites at once (All PSU). Motif search of PUS7 binding motif (f). Identification of UGUAR PUS7 binding motif across newly identified φ sites within KSHV transcriptome.

### Validation of specific Ψ sites in the KSHV transcriptome

Following the identification of Ψ sites by RBS-Seq, validation of a subset of targets was performed to confirm the robustness of the dataset. Initially, RNA immunoprecipitations (RIP) were performed in reactivated TREx BCBL1-Rta cells using a Ψ-specific antibody to precipitate KSHV-encoded RNAs with RBS-Seq mapped Ψ sites (Fig. 5a). This included the PAN and ORF4 RNAs, 28S as a control for a highly pseudouridylated RNA transcript and MBP as negative control previously shown to contain no Ψ sites. Results showed that both PAN and ORF4 RNAs were significantly enriched over MBP, confirming they contain Ψ sites.

**Figure 5.**
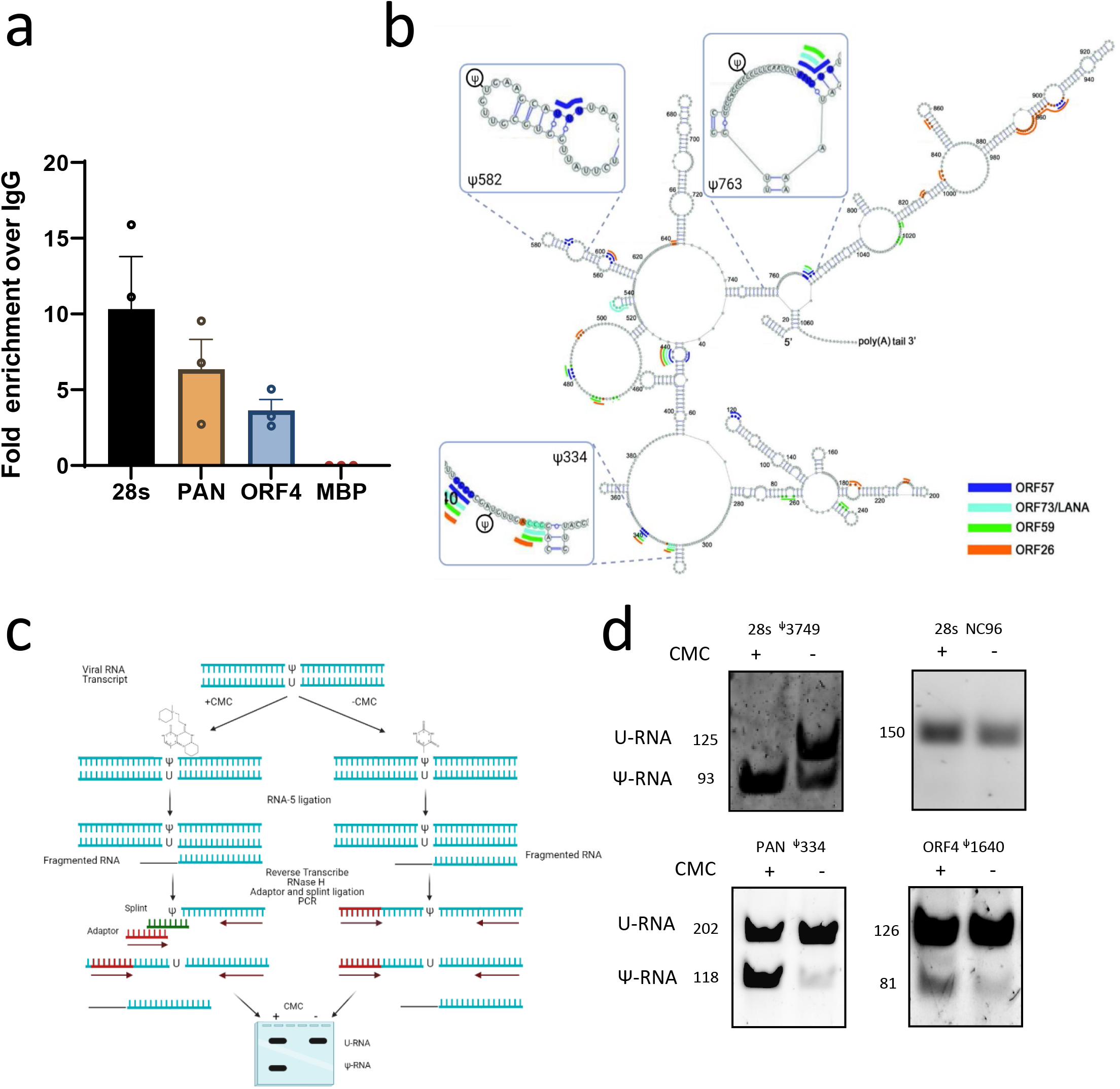
KSHV PAN RNA is pseudouridylated in close proximity to known interaction motifs. RIP analysis of φ modified KSHV genes (a). RIP of TREx BCBL1-Rta cells at 24 hours post reactivation using anti-φ antibody. Followed by qPCR of PAN and ORF4 viral RNAs, with endogenous 28S positive control and negative control MBP transcripts (n = 3). RBS-Seq identified KSHV non-coding RNA PAN φ sites (b). Adapted from Sztuba-Solinska et al 2017 PAN SHAPE analysis, RBS-Seq identified φ sites shown. CMC-ligation assisted PCR (CLAP) schematic (c). Purified RNA from reactivated TREx BCBL1-Rta cells is treated with +/- CMC followed by ligation of an RNA-5 blocker molecule (signified by a black line) to the 3’ end of the fragmented RNA preventing unwanted splint ligation binding. RNA is then reverse transcribed, RNAse H treated and both adaptor and splint ligated to the resulting cDNA. This cDNA is then used as a template for a PCR reaction allowing amplification of both U and φ modified fragments before analysis on a polyacrylamide gel. PAN φ sites confirmed via CMT-RT and ligation assisted PCR analysis (CLAP) (d). CLAP confirming φ site 334 on PAN RNA and site 1640 on ORF4. Ribosmal RNA 28s site 3749 and site 96 used as a positive control and negative control respectively. TREx BCBL1-Rta, PUS1ko or PUS7ko cells were induced for 24 hours before subsequently harvested. RNA was extracted and CLAP performed. Representative image used of 3 repeats.

PAN RNA undertakes multiple essential functions in KSHV lytic replication, specifically acting as a protein scaffold for early expressed viral proteins resulting in enhanced late gene expression ^24, 25^. Therefore, to determine the functional significance of Ψ site modification of the PAN RNA, we first mapped the three Ψ sites identified by RBS-Seq onto a previously identified SHAPE structure of PAN RNA (Fig. 5b) ^26^. These sites, particularly site 334, were noted to be in close proximity to known binding sites of a multitude of viral proteins, including the ORF57 protein. This close proximity indicated that the Ψ site may be involved in the stabilisation of the PAN molecule allowing or disrupting binding of these important viral interactors ^27^. To investigate this possibility we first confirmed that the PAN site 334 was the exact position of the Ψ site, using the recently developed technique, CMC ligation assisted PCR (CLAP) (Fig. 5c) ^28, 29^. CLAP relies on the addition of a CMC adjunct to the Ψ residue. This bulky CMC modified Ψ site acts as a reverse transcriptase terminator thus resulting in shortened DNA fragments reverse transcribed from Ψ modified RNA. Through the addition of both “splint” and “adaptor” short DNA sequences, the CMC fragment can be amplified using the same primers as the non-CMC treated full length fragment. This allows direct semi-quantitative amplification of each fragment and thus relative levels of pseudouridylation can be determined. CLAP was performed on RNA isolated from reactivated TREx BCBL1-Rta cells and the amplification of a shortened transcript confirms that both ORF4 site 1640 and PAN site 334 undergoes Ψ modification (Fig. 5d). Together, RIP and CLAP analysis confirm that the KSHV PAN transcriptome is Ψ modified during lytic replication.

### PUS1/PUS7 knockouts affect PAN stability and poly(A) tail length

The ability of Ψ to influence the stability of mRNA has been previously studied ^30^. Thus, we sought to investigate the effect of PUS1 and PUS7 knockouts on PAN stability. Through utilisation of an Actinomycin D stability assay, we observed in PUS1ko TREx-BCBL1-Rta cells that PAN had a substantial reduction in RNA stability at both 4 and 8 hours post drug treatment and PUS7ko had a reduction primarily at 4 hours post treatment with both knockouts showing decreased half-life of the RNA using an exponential decay model (Fig. 6a). At present the mechanism by which Ψ affects RNA stability is not fully elucidated. Considering the importance of the poly(A) tail on PAN stability, we wished to assess if the loss of stability observed in PUS1 and PUS7ko cells was due to a disruption in the adenylation or deadenylation of the poly(A) tail. G/I extension followed by cDNA synthesis and PCR revealed that while levels of hyperadenylated PAN remained consistent between scrambled control, PUS1ko and PUS7ko (Fig. 6b), there was a significant increase in levels of shorter poly(A) transcripts in both Pus knockout cell lines (Fig. 6c), whereas no significant changes were observed to the poly(A) status of the cellular control gene GAPDH (Fig. 6d). The reduction in poly(A) length within PAN RNA species indicates a possible mechanism for the observed reduction in PAN stability and downstream reduction in expression levels with PUS1 and PUS7 knockout TREx-BCBL1-Rta cells.

**Figure 6.**
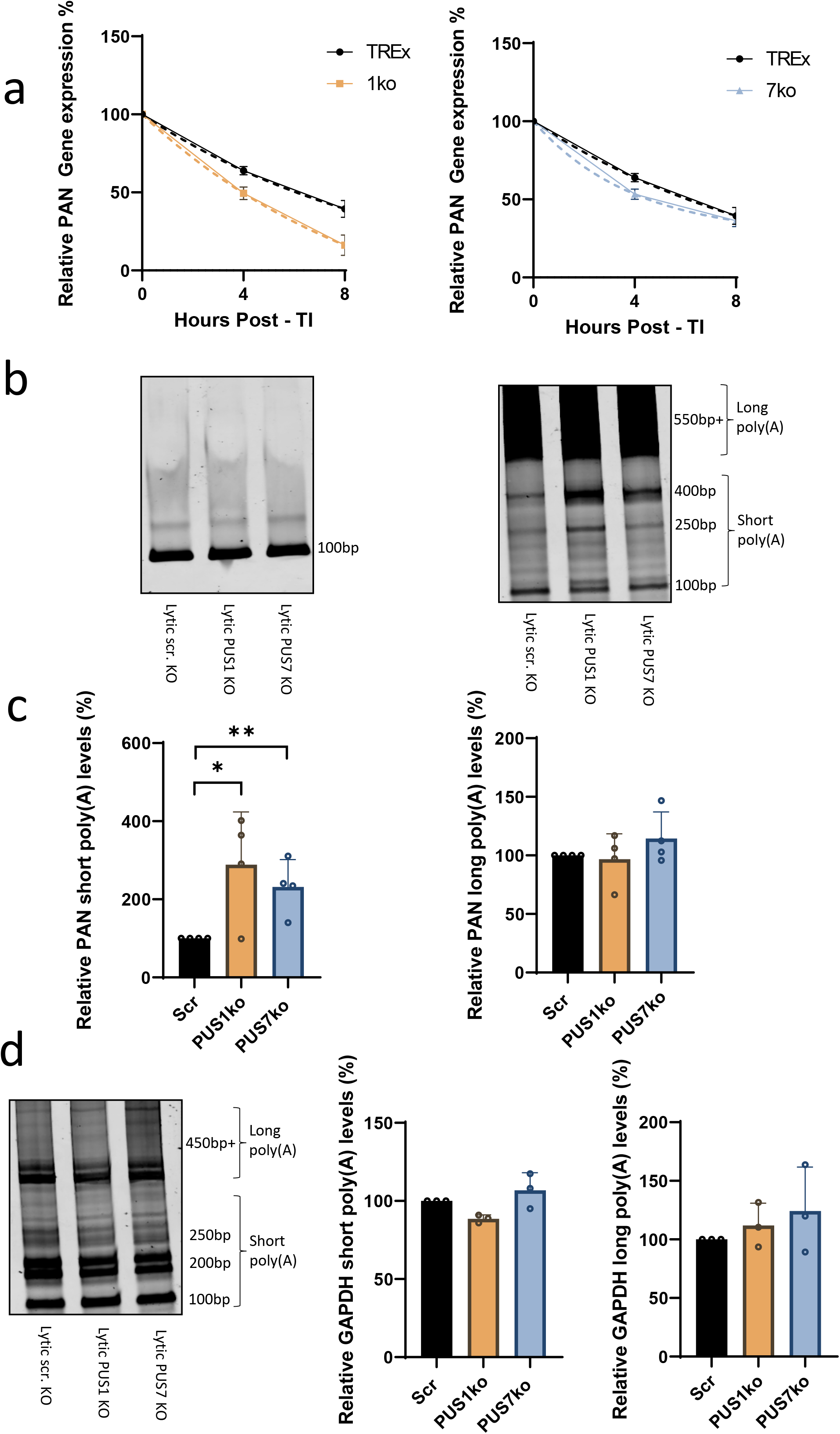
PUS1 and PUS7 knockouts affect PAN poly(A) tail length and stability. The stability of PAN RNA during KSHV lytic reactivation was determined by assessing mRNA decay through Actinomycin D (AcD) treatment of TREx-BCBL1-Rta cells (a). Cells were reactivated using doxycycline 24 hours prior to the addition of 2.5 μg / ml AcD. Cells were then collected at 0, 4 and 8 hours post transcription inhibition (Post-TI) by AcD treatment and total RNA was extracted followed by qRT-PCR. Values were first normalised to GAPDH before normalising to 0 hour time point. An exponential decay model of non-linear regression was performed and plotted. Error bars represent SD, n = 4 for all experiments. poly(A) tail length was assessed through G/I extension followed by cDNA synthesis and PCR (b). TREx-BCBL1-Rta cells were reactivated using doxycycline for 24 hours before harvesting, followed by G/I extension and cDNA synthesis. PCR products of PAN gene specific PCR amplification and PAN poly(A) tail PCR amplification of scrambled (Scr) PUS1 or PUS7 knockout cells were analysed by acrylamide gel electrophoresis using a Li-COR Odyssey SA imager. Representative images of n = 4 shown. Densitometry analysis of long poly(A) and short poly(A) acrylamide gels (c). Gene specific and poly(A) amplification of GAPDH and corresponding densitometry (d). Values were normalised to densitometry of gene specific PCR product before normalisation to Scr. Error bars represent SD, n = 4 for all experiments. p ≤ 0.05 *, p ≤ 0.01 ** using a two tailed Students unpaired t-test.

### PAN mutant 334 shows important function for PAN stability and expression

Knockout of PUS1 or PUS7 has been shown to result in global reduction of pseudouridylation at PUS1 or PUS7 sites respectively, therefore the effects on PAN stability observed in PUS1 and PUS7 knockouts cannot be directly attributed Ψ within PAN. Thus, to isolate Ψ function in PAN RNA from other KSHV Ψ modifications and their effects, we performed a CRISPR cas9 knockout of PUS1 and PUS7 in HEK-293T cells (Fig. 7a). ORF57, an important viral protein involved in a number of essential viral processes has previously been shown to significantly enhance PAN expression ^27, 31^. To explore the functional implications of Ψ modification on PAN and its relation to ORF57, we next determined if the ORF57 protein was able to enhance PAN RNA levels in PUS1 or PUS7 knockout cells (Fig. 7b). Interestingly, both PUS1ko and PUS7ko showed a significant reduction in the enhancing effect of ORF57 on PAN expression. Furthermore, due to the close proximity of the Ψ site to local ORF57 binding sites, we wanted to assess if binding of the ORF57 protein to PAN RNA could be affected upon loss of PUS enzyme activity. Initially, we co-transfected PAN with ORF57-eGFP in our control HEK-293T or PUSko HEK-293T cell lines and carried out a RNA immunoprecipitation using GFP-TRAP beads pulldown and bound PAN RNA levels were assessed (Fig. 7c-d). PUS1ko and PUS7ko resulted in reduced binding of PAN to ORF57-eGFP, while pre-BTG1 positive control levels remained constant suggesting the presence of Ψ on PAN is important for ORF57 binding allowing enhanced expression.

**Figure 7.**
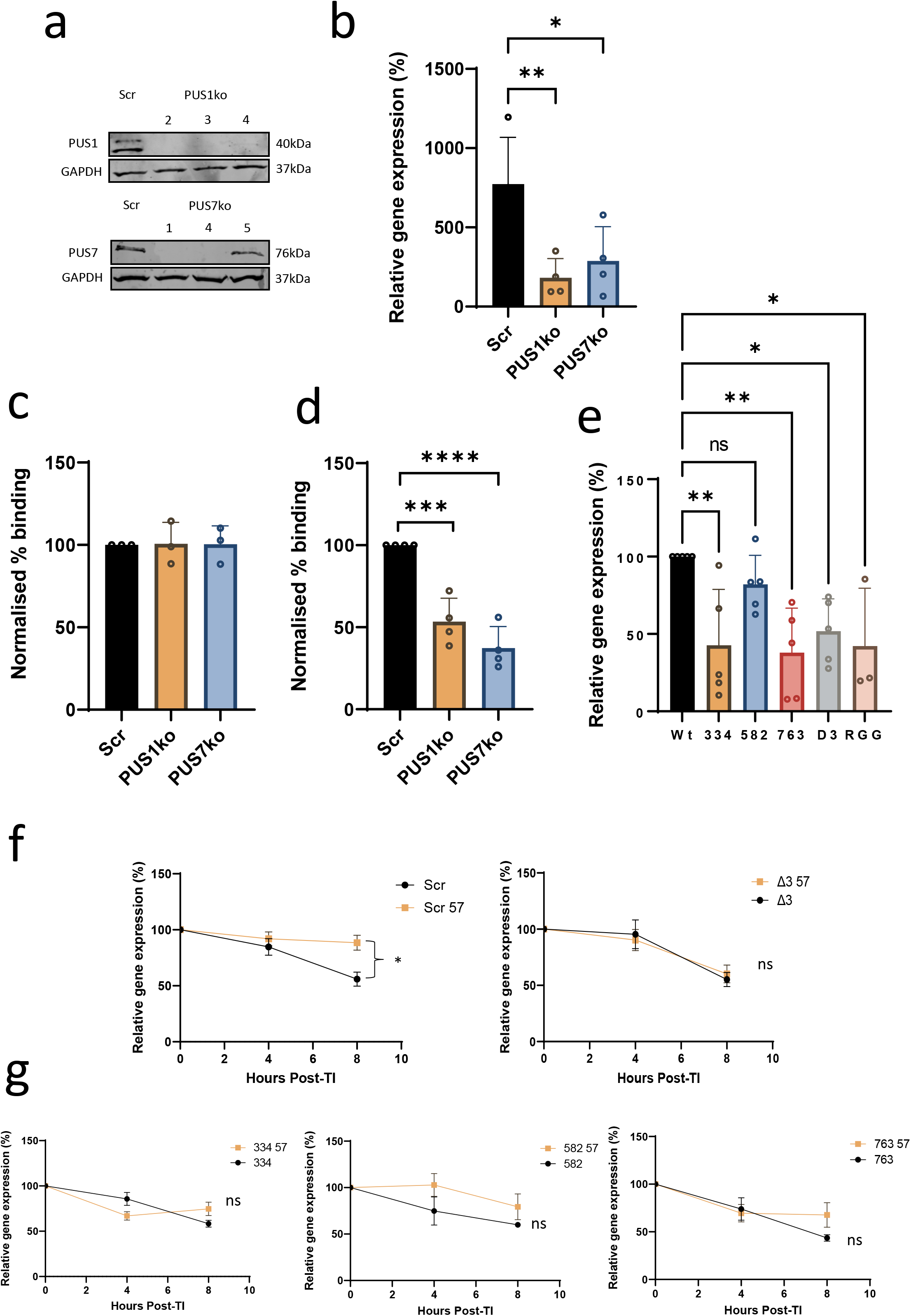
φ site 334 is important for ORF57 enhancement of PAN expression and directly affects PAN stability. Knockout of PUS1 and PUS7 in HEK-293T cells as confirmed via western blotting (a). Overexpression of PAN and ORF57 in PUS1ko and PUS7ko HEK-293T cells (b). HEK-293T cells were transfected with pCMV-PAN alone or pCMV-PAN with pCMV-ORF57-eGFP before harvesting after 48 hours. Samples were first normalised to GAPDH before normalising to respective PAN only sample. RIP analysis of ORF57-eGFP in HEK-293T cells. HEK-293T cells were transfected with pCMV-PAN and pCMV-ORF57-eGFP and an RIP was performed at 24 hours post transfection using GFP-TRAP beads. Followed by qPCR of PAN (c) or pre-BTG1 (d) genes and GAPDH (n ≥ 3). RIPs are expressed as fold change over GAPDH following normalisation to scrambled control. Overexpression of PAN pseudouridine mutants co-transfected with ORF57-eGFP (e). HEK-293T cells were transfected with pCMV-PAN (or mutant pCMV-PAN) or both pCMV-PAN (or mutant pCMV-PAN) and pCMV-ORF57-eGFP before harvesting after 48 hours. Samples were RNA extracted and analysed via qRT-PCR. Samples were first normalised to GAPDH before normalising to respective PAN only samples and WT PAN. Error bars represent SE, n = 4 for all experiments, p ≤ 0.05 *, p ≤ 0.01 ** using a one way ANOVA with Dunnett post-test. RNA stability of WT PAN and Δ3 Ψ PAN mutants (f) or 334, 582 and 763 mutants (g) co-transfected with ORF57-eGFP. The stability of PAN RNA or φ negative mutant transcripts were determined by assessing mRNA decay through AcD treatment of transfected HEK-293T cells. Cells were transfected 24 hours prior to the addition of 10 μg/ml AcD. Cells were then collected at 0, 4 and 8 hours post transcription inhibition (Post-TI) by AcD treatment and total RNA was extracted followed by qRT-PCR. Error bars represent SE, n = ≥ 5 for all experiments, p ≤ 0.05 *, p ≤ 0.01 ** using a two tailed Students unpaired t-test.

To further examine the specific Ψ sites within PAN and how they contribute to PAN function, residues identified as Ψ sites with RBS-Seq, namely 334, 582 and 763 were mutated individually or as a triple mutant (Δ3) containing all three mutations. A T-A substitution was chosen to maintain GC content and to minimise any effects to overall RNA structure. To assess the effect of mutating individual Ψ sites within the PAN RNA, co-transfection assays were performed for each individual mutant in the presence of a KSHV ORF57 expression construct (Fig. 7e) ^32^. Assessing the expression levels of each PAN mutant in the presence of ORF57, results determined that mutant 334 and the Δ3 mutant showed significantly reduced PAN expression levels in comparison with WT PAN RNA. These levels were comparable with the negative ORF57 RNA binding mutant, RGG ^33^, which showed no enhancement of PAN RNA levels. These results implied that the 334 and Δ3 mutants may affect the stability of PAN RNA. Therefore to assess the stability of the PAN Ψ mutants an Actinomycin D stability assay was performed in the absence or presence of the ORF57 protein (Fig. 7f-g). As expected, the stability enhancing effect of ORF57 for both mutant 334 and Δ3 was ablated. While non-significant, both mutant 582 and 763 stability appeared to be similar to wildtype PAN RNA levels in the presence of ORF57 protein, highlighting that the Ψ site at nucleotide 334 is key in enhancing PAN RNA stability in the presence of the ORF57 protein. Additionally, the stability of the PAN RNA mutants in the absence of ORF57 was unaffected, indicating that the mutation alone is insufficient to cause decreased stability thus showing an ORF57 specific stabilising phenotype (Supplementary Fig. 4). Together these data show Ψ is important for ORF57-mediated functional enhancement of PAN expression and the stability of PAN provided by ORF57 in the presence of specific Ψ sites is likely important for this enhancement.

## Discussion

This study is the first to investigate the transcriptome-wide role of Ψ within a DNA virus. KSHV was used as a model pathogen since it is known to manipulate and utilise a large number of host cell proteins and pathways during its replication cycle, including other RNA modifications ^18^. Redistribution of PUS7 to KSHV RTCs during lytic reactivation reinforced the hypothesis that KSHV may utilise pseudouridylation to positively modify viral RNA transcripts. These virus-induced intranuclear structures, enable multiple processes required for KSHV lytic replication to occur, including viral transcription, viral DNA synthesis and capsid assembly. By undertaking a quantitative proteomic approach to identify PUS enzyme interaction partners, we showed a comprehensive interactome of cellular proteins that drastically change under viral stress, thus opening further avenues of research for their important role in both cellular function but also their involvement in viral protein-protein or protein-RNA interactions. Notably, PUS7 redistribution to KSHV RTCs leads to a dramatic change in the PUS7 interactome, with a large number of host protein-protein interactions found during KSHV latency no longer occurring upon reactivation into the lytic replication phase. Interestingly, PUS7 predominantly loses interactions with histones H2B H1.4 and H4 that have previously been confirmed as associated proteins through ChIP-MS analysis ^34^. ChIP-qPCR for PUS7 probing for enhancers and promoters showed a significant enrichment highlighting PUS7 role in co-transcriptional loading. Furthermore, interactions with heterogeneous nuclear ribonucleoproteins hnRNP C1/C2, hnRNP A2/B1 and hnRNP U are also lost. hnRNP C has previously been shown to contain binding site that overlaps intronic Ψ ^7^. The loss of such interactions during KSHV lytic replication may influence pre-mRNA splicing, mRNA-protein interactions or modifying pre-mRNA structure and stability.

Surprisingly, only a small number of interactions increase, including a number of viral proteins such as ORF59 and the cellular stress protein HSP90. This sequestering into KSHV RTCs may serve the virus twofold; the utilisation of PUS7 pseudouridylation activity for KSHV lytic transcripts such as PAN and ORF4 occurring co-transcriptionally, and/or the reduction of Ψ on cellular genes. While PUS1 maintains a generalized localisation around both the nucleus and cytoplasm, there is still a distinct change in protein-protein interactions upon viral reactivation. These interactions are predominantly increased in cell stress proteins and translational factors along with viral proteins. In particular, PUS1 showed an increased interaction with HSP90 during KSHV lytic replication, which has previously been shown to increase PUS7 stability and expression ^35^. Additionally, PUS1 interaction with EEF1A1 and HSP60 are increased, both of which mRNAs contain experimentally verified Ψ sites that are modified under stress conditions ^36, 37^, suggesting the interaction of PUS1 with these proteins may be stress induced. One of PUS1 primary pseudouridylation targets is tRNA ^38^, which may have a significant impact in protein translation, suggesting this function may be modified during KSHV lytic reactivation. It must also be noted that an important aspect of KSHV lytic reactivation is host-cell shutoff, involving KSHV SOX-mediated degradation of cellular RNAs ^39, 40^, and as Ψ has been shown to directly affect the stability of RNA molecules, it is possible that viral changes to the PUS1 and PUS7 interactomes could affect overall stability of a number of cellular genes, contributing to the overall host cell shutoff of genes not essential for the virus. PUS1s localisation and interaction with numerous KSHV proteins suggests PUS1 may carry out pseudouridylation of viral transcripts in both the cytoplasm and nucleus. Conversely, PUS7 interaction profile suggests that localisation to viral RTCs reduces interaction with cellular proteins and indicates PUS7 may be involved in co-transcriptionally pseudouridylating viral RNA transcripts (Fig. 2d).

Through the use of CRISPR cas9 knockouts, we show that both PUS1 and PUS7 are essential for KSHV lytic replication. The ~30% reduction in expression of the immediate-early PAN RNA and early ORF57 RNA and protein, compared to the significant 80% reduction of late ORF65 protein in both PUS1ko and PUS7ko cells indicates that Ψ is not directly involved in the latent/lytic switch but more likely affecting one or more viral processes that occur downstream in the lytic temporal cascade. To further understand the effect of the PUS enzymes on viral replication, we sought to determine if the reduction in Ψ affect the virus was purely changes to the cellular transcriptome or if the virus transcriptome itself was Ψ modified. It has been previously shown through transcriptome wide analysis that the human pathogens *Trypanosoma Brucei,* Influenza A and HIV have Ψ modified transcripts ^30, 41, 42^ however there is limited research on the functional role of Ψ in the context of viral infection. Our RBS-Seq experiment revealed that the KSHV transcriptome is heavily Ψ modified. By performing RBS-Seq at 8 and 20 hours post reactivation, we wished to determine if any modifications were changed as the lytic temporal cascade progresses and as cell stress intensifies. However, modifications that were present at 8 hours post reactivation were mostly consistent at 20 hours post reactivation, providing evidence that the Ψ modification may be dynamically induced throughout reactivation ^43^. Mapping the Ψ sites to genome features revealed the distribution of Ψ to be preferentially found on the CDS of the KSHV transcriptome. While the majority of traditional Ψ found in human or yeast cellular transcripts are also within the CDS, there are proportionally more sites found in UTR regions, particularly 3’ UTR than within the KSHV transcriptome ^23^. Examining Ψ sites in alternative topological features showed a significant number of sites in alternative start site regions which may have implications on downstream protein expression (Supplementary Fig. 5). Additionally there is a high proportion of Ψ deletions identified in the repeating regions of KSHV. This is most likely an artifact of the deletion mechanism of the DNA reverse transcriptase which shows increased deletions at repeating regions out with Ψ status. Motif analysis of nucleotides surrounding Ψ sites revealed differential motifs between the CDS, intergenic regions and all sites analysed together. Recent research has shown that motif binding alone is not sufficient to predict Ψ sites, however these motifs likely represent different proportional pseudouridylation by all PUS enzymes. When analysing all Ψ sites, the AGGA and AAAA motifs show it is likely that PUS1s HRU binding motif would contribute a significant proportion to the motif analysis and thus, likely to pseudouridylate the most Ψ sites. Comparatively, PUS7s specific binding motif was found in just 7 Ψ sites. While out with the scope of this study, the RBS-Seq dataset can be further examined to identify Ψ within cellular genes and any changes that may occur during KSHV lytic reactivation that could influence the overall landscape of anti-viral or pro-viral cellular genes.

Validation of the non-coding viral PAN and ORF4 RNAs was performed via an RNA-immunoprecipitation using a Ψ-specific antibody to immunoprecipitate Ψ modified RNA. PAN and ORF4 were selected as two high confidence hits from RBS-Seq and also for their importance for viral replication ^44^. Furthermore, PAN was selected for further study due to its huge abundance during viral reactivation, accounting for >90% of viral reads within a cell, and thus we surmised that PAN may lead to significant changes in virus replication in the absence of Ψ. There has been growing evidence that Ψ can directly affect levels of protein translation, ^11, 45^ thus we wished to examine further the lesser known effects of Ψ on a viral non-coding RNA. Within PAN, three Ψ sites were detected at nucleotides 334, 582 and 763. Due to the proximal nature of multiple viral protein binding sites surrounding site 334, we hypothesised site 334 may be important in influencing binding of viral proteins essential for viral replication ^26^.

When assessing the expression levels of PAN in our PUS1ko and PUS7ko TREx BCBL1-RTA cells during lytic reactivation, we noted a significant reduction in PAN expression within both PUS1ko and PUS7ko cells. The reduction of this immediate-early gene and PANs functional significance on late gene expression highlights Ψ drastic effect on KSHV replication. While the roles of varying poly(A) tail lengths in mRNA is currently broad and often undefined ^46^, we identified a possible source of reduced PAN expression through an increase in the pool of short poly(A) tails found within both PUS1 and PUS7ko cells. Coupled with the reduced stability of PAN during reactivation in PUS1 and PUS7ko cells, and a larger reduction in stability in PUS1ko corresponding with a larger change in poly(A) tail length, this suggests one possible mechanism of how Ψ may directly affect PANs function.

During reactivation, there are thousands of Ψ modifications occurring in both cellular and viral transcripts and thus focusing on single transcript modifications with full PUS knockouts is a significant challenge. To isolate PAN from the multitude of associating viral factors during lytic reactivation, we generated PUS1 and PUS7ko HEK-293T cells with which we could assess PAN function through transfection. Previous work has shown that PAN has a number of interaction motifs enabling ORF57 protein recruitment, including an Mta responsive element (MRE) and expression and nuclear retention element (ENE). These regions are important for achieving high levels of expression, up to 20 fold increase and also nuclear retention ^31^. Here we show that the expression enhancing effect of ORF57 is impaired in both PUS1 and PUS7ko HEK-293T cells, consistent with the reduction seen in TREx BCBL1-RTA PUSko cells. Ψ has been shown to reduce RNA-protein binding affinity when the modification is located directly within the binding motif ^8^ however in this case, the 334 modification is 4-6 bases from the sites and sites 582 and 763 were 7 and 13 bases away respectively. Interestingly, PUS1ko and PUS7ko both reduce the overall binding of ORF57 to PAN, indicating that the modification is likely affecting the binding sites through changes to secondary structure rather than immediate changes to binding dynamics within the motif. Additionally, these sites are not located in the MRE or ENE elements that have been shown to improve PAN stability. However, this impaired binding is likely a contributor to the reduction in PAN expression highlighting the importance of all multiple binding regions on PAN. Further work may identify Ψ importance in the impaired binding of other important viral factors such as ORF59 or cellular factors such as PABPC1 and Aly/REF ^31, 47^ and whether this is impaired binding directly affects the polyadenylation of PAN.

Single Ψ sites can affect the dynamics of an RNA molecule. By mutating each Ψ PAN site individually, and generating a Δ3 Ψ PAN mutant, we showed that both ΔPAN334 and Δ3 mutants followed a phenotype of reduced PAN expression enhancement by ORF57. Interestingly, when examined further, ORF57 functioning as a stabiliser of PAN is also reduced in both ΔPAN334 and Δ3 mutants. This shows that PAN 334 is a Ψ site potentially important for allowing the interaction between PAN and ORF57, stabilising PAN and allowing enhanced PAN expression. Conversely, the landscape of Ψ is broad as to affect many cellular and viral genes ^36^, of which there can be multiple interactions affect PAN molecular dynamics. Thus, while clearly essential for PAN function, Ψ knockdown can affect numerous RNAs across the host and virus transcriptome making examining direct function challenging. Further work making use of RBS-Seq with PUS knockouts in both latent and lytic KSHV reactivation, examining both the viral and host transcriptome, would allow a more broad overview of how the landscape of Ψ changes, and may allow more elucidation of Ψ function with KSHV replication.

## Methods

### Reagents tables

Plasmids (Supplementary Table 1, antibodies (Supplementary Table 2) and primers (Supplementary Table 3).

### Cells lines and reagents

TREx BCBL1-Rta cells are a genetically engineered BCBL-1 primary effusion lymphoma (PEL) B cell line that expresses Myc-tagged RTA under a doxycycline inducible promoter, a gift from Professor Jae U. Jung (University of Southern California, USA). TREx BCBL1-Rta cells were cultured in RPMI1640 growth media with glutamine (Gibco®) supplemented with 10% (v/v) fetal bovine serum (FBS, Gibco®), 1% (v/v) penicillin-streptomycin (P/S, Gibco®) and 100 μg/ul hygromycin B (Thermo Scientific). For virus reactivation, RTA expression was induced through the addition of 2 μg/mL doxycycline hyclate (Sigma-Aldrich). HEK-293Ts were purchased from ATCC (American Type Culture Collection) and cultured in Dulbecco’s modified Eagle’s medium with glutamine (DMEM, Lonza) supplemented with 10% (v/v) FBS and 1% P/S. The antibodies used throughout this study include anti-ORF57 (Santa Cruz, sc-135746 1:1,000), anti-ORF65, anti-GAPDH (Abcam, ab8245 1:5,000), anti-PUS1 (SIGMA-ALDRICH, SAB1411457 1:1000) and anti-PUS7 (Invitrogen™, PA5-54983, 1:1000).

### CRISPR stable cell lines

HEK-293T cells were transfected with the 3 plasmid lentiCRISPR v2 system. In 12-well plates, 4 ul of lipofectamine 2000 (Invitrogen™) was combined with 1 ug lenti CRISPR V2 plasmid (a gift from Feng Zhang, Addgene plasmid #52961) expressing the guide RNA (gRNA) targeting the protein of interest, 0.65 μg of pVSV.G and 0.65 μg psPAX2. pVSV.G and psPAX2 were gifts from Dr. Edwin Chen at the University of Leeds. Two days post transfection the viral supernatant was harvested, filtered (0.45 um pore, Merck Millipore) and used to transduce TREx BCBL1-Rta cells in the presence of 8 μg/mL of polybrene (Merck Millipore). Virus supernatant was removed 6 hours post transduction and cells were maintained for 48 hours before puromycin (Sigma-Aldrich) selection. Stable mixed population cell lines were maintained until confluent before single cell selection. Single cell populations were generated through serial dilution of ~100 cells in 96 well plates. Positive wells were cultured for 3-5 weeks and maintained with fresh media before transferal into 6 well plates. Upon confluence, clones were tested via western blot for expression of target protein of interest.

### Viral reinfection assay

TREx BCBL1-Rta, TREx BCBL-Rta PUS1ko or PUS7ko cells were reactivated and harvested after 72 h as previously described ^48^. Cellular supernatant was filtered with a 0.45 μm pore filter (Merck Millipore) and subsequently used to inoculate HEK-293T cells at a 1:1 ratio with DMEM tissue culture media. Active KSHV transcription was quantified at 48 h post-infection by RT qRT-PCR. Total RNA was extracted from cell lysates using RNeasy Mini Kit as described by the manufacturer. cDNA synthesis was carried out with 1 μg total RNA using LunaScript™ RT SuperMix Kit according to the manufacturers protocol. Subsequent qPCR reactions were carried out using ORF57 and GAPDH specific primers as described in the qRT-PCR method.

### Viral episome count assay

TREx BCBL1-Rta, TREx BCBL-Rta PUS1ko or PUS7ko cells were serially passaged and cells harvested after 14 days. Total DNA was extracted from cell pellets using Monarch Genomic DNA Purification Kits (New England Biolabs) and viral episome copies quantified by qPCR of the viral gene ORF57 as described in the qRT-PCR method.

### Two-step reverse transcription quantitative PCR (qRT-PCR

Total RNA from cell pellets was extracted using a Monarch Total RNA Miniprep kit (New England Biolabs) according to the manufacturer’s instructions. Reverse transcription was performed on 500 ng of total RNA using a LunaScript™ RT SuperMix Kit (New England Biolabs) as according to the manufacturer’s instructions. Quantitative PCR (qPCR) reactions included 10 μl 1 X GoTaq® qPCR Master Mix (Promega), 0.5 μM of each primer and 5 μl template cDNA. Cycling was performed in a RotorGene Q 2plex machine (Qiagen). The cycling programme used was; a 10 minute initial preincubation at 95 °C, followed by 40 cycles of 95 °C for 15 s, 60 °C for 30 s and 72 °C for 20 s. A melt curve step was performed post qPCR to confirm single product amplification. Gene expression analysis was performed with normalisation to the housekeeping gene GAPDH (ΔC_T_) and reference sample (ΔΔC_T_).

### RNA stability assay

TREx BCBL1-Rta cells were reactivated with doxycycline hyclate. At 24 hours post reactivation, cells were treated with 2.5 μg/ml of actinomycin D (Thermo Scientific) and samples were collected at 0, 4 and 8 hours post treatment. HEK-293T cells were transfected with 500 ng pCMV-PAN or pCMV-PAN mutant and 500 ng pCMV-ORF57-eGFP. At 24 hours post transfection, cells were treated with 10 μg/ml of actinomycin D. Total RNA was extracted using Monarch® Total RNA Miniprep Kits (New England Biolabs) according to the manufacturer instructions. cDNA synthesis was carried out using LunaScript™ RT SuperMix Kit. qRT-PCR was performed as described above. Normalisation was carried out using GAPDH and data was further normalised to 0 hour sample. The non-linear regression analysis was applied to a one phase decay model as allowed by decay parameters in TREx-BCBL1-Rta stability assays. One phase decay model used in Graphpad Prism 9 (GraphPad Software, www.graphpad.com) as Y = (Y0 – Plateau) * exp (-K * X) + Plateau, were Y0 is 100%, K is the rate constant with X as minutes. Tau is time constant as the reciprocal of K.

### Poly(A) Tail-Length Assay

Scrambled or PUS1/PUS7ko TREx BCBL1-Rta cells were reactivated with doxycycline hyclate and harvested at 24 hours post reactivation. Total RNA was extracted using Monarch® Total RNA Miniprep Kits (New England Biolabs) as described by the manufacturer. Total RNA was G/I tailed, reverse transcribed and underwent PCR using a poly(A) Tail Length Assay Kit (Invitrogen™) according to manufacturer’s instructions. PCR was performed using forward and reverse gene specific primers, or gene specific primers and poly(A) universal reverse primer for both PAN and GAPDH. Samples were then loaded onto an 8% polyacrylamide gel alongside a 50 bp ladder (New England Biolabs) and resolved at 100 V for 45 minutes in TBE buffer. The gel was stained for 20 minutes with 1:10,000 SYTO™ 60 stain (Invitrogen™) before subsequent visualisation on a Odyssey® CLx (LI-COR). Densitometry analysis was performed using Image Studio™ (LI-COR).

### RNA immunoprecipitations

For Ψ RIPs, TREx BCBL1-Rta cells were reactivated with doxycycline hyclate and harvested 24 hours post reactivation. Cells were lysed and RNA extracted with TRIzol LS (Invitrogen™) as per manufacturer’s instructions. 10 μg total RNA was incubated overnight at 4 °C with Dynabeads™ Protein G magnetic beads pre-bound with anti-Ψ (Diagenode) according to manufacturer’s instructions. Following pulldown, RNA samples were incubated with Proteinase K buffer (10 mM Tris pH 7.5, 150 mM NaCl, 0.5 mM EDTA, 10% SDS, proteinase K) for 30 minutes at 55 °C minutes before further RNA extraction with TRIzol LS (Invitrogen™). cDNA was synthsised using using LunaScript™ RT SuperMix Kit (New England Biolabs) before analysis via qRT-PCR. Samples were analysed using fold enrichment over GAPDH before further normalisation with scrambled samples.

For GFP RIPs, scrambled or PUS1/PUS7ko HEK-293Ts were transfected with 2 μg PAN and 2 μg ORF57-eGFP plasmids using Lipofectamine 2000 (Thermo Fisher Scientific) according to manufacturer’s instructions. Cells were lysed before incubation with GFP-Trap Agarose beads (Chromotek) overnight at 4 °C using manufacturer’s instructions. Samples were incubated with Proteinase K buffer for 30 minutes at 55 °C before RNA extraction with TRIzol LS (Invitrogen™) as per manufacturer’s instructions. cDNA was synthesised using LunaScript™ RT SuperMix Kit (New England Biolabs) before analysis via qRT-PCR. Samples were analysed using fold enrichment over GAPDH before further normalisation with scrambled samples.

### Quantitative proteomics

Protein A beads pre bound with anti-PUS1, anti-PUS7 or IgG control antibodies were incubated with latent or reactivated TREx BCBL1-Rta cell lysate. Immunoprecipitated samples were sent to the University of Bristol Proteomics facility for TMT LC-MS/MS. A detailed protocol was followed as previously described ^20, 49^. Briefly, samples were trypsin digested and labelled with amine-specific isobaric tags resulting in differentially labelled peptides of the same mass. Labelled samples were pooled and fractionated using Strong Anion eXchange chromatography before analysis by synchronous precursor selection MS3 on an Orbitrap Fusion Tribrid mass spectrometer (Thermofisher) controlled by Xcalibur 2.1 software (Thermo Scientific) and operated in data-dependent acquisition mode. Raw data files were processed and quantified using Proteome Discoverer software v1.4 (Thermo Scientific) compared against UniProt Human database (downloaded October 2019) plus KSHV protein sequences using SEQUEST algorithm.

Data was first analysed by the removal of background abundance values from PUS1 and PUS7 abundance values. Background values were generated from the TREx BCBL1-Rta IgG control pulldown. A cutoff abundance value of 100 was selected as a minimum detection level for further analysis. The mass spectrometry proteomics data have been deposited to the ProteomeXchange Consortium via the PRIDE partner repository with the dataset identifier PXD037379 ^50, 51^.

### Immunoblotting

Protein samples were run on 12% polyacrylamide gels and transferred to nitrocellulose membranes (GE Healthcare) via semi-dry transfer in a Bio-Rad Trans-blot Turbo transfer machine. Membranes were blocked with TBS + 0.1% (v/v) Tween 20 (TBS-T) and 5% (w/v) dried milk powder (Marvel) for 1 hour. The membrane was then incubated with relevant primary, followed by secondary antibodies for 1 hour incubations diluted in 5% (w/v) milk TBS-T. Membranes incubated with secondary fluorescent antibodies were dried and imaged on an Odyssey® CLx (LI-COR).

### Immunofluorescence

Sterilised glass coverslips were treated with Poly-L-Lysine for 15 mins before seeding TREx BCBL1-Rta cells. After 8 hours post seeding, TREx cells were reactivated with doxycycline hyclate. Cells were fixed at 24, 48 or 72 hours post reactivation with 4% (v/v) paraformaldehyde in PBS for 15 minutes. Subsequently, wells were washed twice in PBS and permeabilised in PBS containing 1% Triton X-100 for 10 minutes. Coverslips were blocked with 5% (v/v) BSA in PBS for 1 hour before subsequent incubation with primary and secondary antibodies, both for 1 hour at 37 °C. Glass coverslips were then mounted onto microscope slides using Vectashield® Hardset with DAPI. Slides were visualised on a Zeiss LSM 880 laser scanning confocal microscope and images analysed using Zen® 2011 (Zeiss).

### CMC-ligation assisted PCR (CLAP)

40 μg of total RNA harvested from TREx BCBL1-RTa cells was CLAP treated as according to Zhang et al 2022 ^29^. In brief, 40 μg of total RNA was divided and 20 μg treated with CMC, 20 μg without, in TEU buffer at 30 °C for 16 hours. The reaction was subsequently stopped with KOAc KCL (Stop buffer) followed by 2x 75% ethanol wash steps with 2 hour −80 °C incubations. Following washes, CMC adjuncts underwent reversal through the addition of 50 mM Na2CO3 supplemented with 2 mM EDTA (Reversal Buffer) and incubated for 6 hours at 37 °C before 1 further 75% ethanol wash 2 hours at −80 °C. Samples then underwent phosphate group addition using T4 PNK at 37 °C for 30 min. Subsequently, samples were ligated with an RNA-5 oligo using T4 ligase 1 incubated at 16 °C for 16 hours. Samples were then reverse transcribed using a target specific RT primer and using AMV transcriptase at 42 °C for 1 hour before a denaturation step at 85 °C for 5 min. Samples were further treated with RNase H at 37 °C for 20 min before denaturation at 85 °C for 5 min. Adaptor and splint oligo’s were combined in a 1:1 ratio to form a oligo/splint mixture that was added to above RT mixture and incubated at 75 °C for 3 min, to which DNA ligase buffer and enzyme plus DMSO were added and incubated at 16 °C for 16 hours, followed by inactivation at 65 °C for 10 min. Samples were subsequently analysed via KAPA2G PCR using the following conditions; Initial denaturation 95 °C for 3 min followed by 10 cycles of 95 °C for 15 s, 65 °C for 15 s (descending 1 °C per cycle) and 72 °C for 5 s, then 10 cycles of 95 °C for 15 s, 55 °C for 15 s and 72 °C for 5 s, with a final extension of 72 °C for 1 min. PCR products were ran on 10% DNA polyacrylamide gels.

### RBS-Seq RNA Isolation and Preparation

TREx BCBL1-Rta cells were seeded at 5×10^5^ cells per ml in T25 flasks. Cells were reactivated with 2 μg / ml doxycycline hyclate and harvested at 8 hours and 20 hours post reactivation. Total RNA was isolated using TRIzol Reagent (Invitrogen) and samples were depleted of rRNA using the NEBNext® rRNA Depletion Kit (Human/Mouse/Rat) as described in the manufacturer’s instructions. Samples were then split, half were untreated and half of which were bisulfite treated using an EZ RNA Methylation Kit (Zymo Research) according to manufacturer’s instructions. Samples were sent to Centre for Genomic Research, Liverpool (United Kingdom) for library preparation.

### Library Preparation and Sequencing

The NEBNext Ultra II Directional RNA Library Prep Kit for Illumina was used to generate paired-end libraries at the University of Liverpool Centre for Genomic Research. In short, Antarctic phosphatase (New England Biolabs) and polynucleotide kinase (PNK) (New England Biolabs) were sequentially applied on the samples. First, for each sample ~75–100 ng of the RNA was diluted in RNase-free water (16 μl total). Then 2 μl of 10× phosphatase buffer, 1 μl of Antarctic phosphatase and 1 μl of RNase inhibitor were added and mix well, followed by 30 min incubation at 37 °C, then 5 min incubation at 65 °C, and then kept on ice. Next, to each sample, 17 μl of nuclease-free water, 5 μl of 10× PNK buffer, 5 μl of 10 mM ATP, 1 μl of RNase inhibitor and 2 μl of PNK were added and mixed well followed by incubation for 60 min at 37°C then kept on ice. The end-repaired RNA was then cleaned up with an RNeasy MinElute cleanup kit (QIAGEN) according to the manufacturer’s instructions and eluted in 14 μl of RNase-free water. For 3’-adapter ligation, 2 μl of the v1.5 sRNA 3’ adapter (Illumina) was mixed with the 14 μl eluate of the previous step in a 200 μl nuclease-free, thin-walled PCR tube, followed by 2 min incubation at 70 °C on a preheated thermal cycler then kept on ice. Next, 2.5 μl of 10x T4 RNA ligase 2 (truncated) reaction buffer, plus 2 μl of 100 mM MgCl_2_, 1 μl of RNase inhibitor and 3 μl of T4 RNA ligase 2 (truncated) (New England Biolabs) were added and mix well by pipetting, followed by 1 h incubation at 22 °C on a preheated thermal cycler, then kept on ice. For 5’-adapter ligation, for each sample 2 μl of 5’ adapter (Illumina) (total 12 μl) was put in a new 200-μl nuclease-free, thin-walled PCR tube and on a preheated thermal cycler, heat-denatured at 70 °C for 2 min, then kept on ice. Then 2 μl of this heat-denatured 5’ adapter was added to each of the 3’-adapter ligation tubes (from previous step), plus 3 μl of 10 mM ATP, and 2 μl of T4 RNA ligase (New England Biolabs) and mixed, followed by incubation for 1 h at 20 °C on a preheated thermal cycler. For first-strand cDNA synthesis, 4 μl of the 3’-5’-adapter-ligated RNA was mixed with barcoded RT primers (Illumina) and cDNA synthesis was performed using SuperScript™ II Reverse Transcriptase (Invitrogen) according to the manufacturer’s instructions. For amplifying the cDNA library, 10 μl of the cDNA was mixed with 10 μl of 5x Phusion high-fidelity buffer, 1 μl of 10mM dNTP mix, 1 μl of forward and 1 μl of reverse 25 μM PCR primers (Illumina), and 0.5 μl of Phusion high-fidelity DNA polymerase (New England Biolabs), then reached to final reaction volume of 50 μl by addition of 26.5 μl of Nuclease-free water. Next on a thermal cycler, the PCR mix was denatured at 98°C for 30 s, followed by 15 cycles of 98 °C (30 s), 60 °C (30 s), 72 °C (15 s), and a final incubation at 72 °C for 10 min, then kept at 4 °C. For library clean up, Agencourt Ampure XP beads (Beckman Coulter) were applied on the amplified libraries according to the manufacturer’s instructions. The resulting libraries were sequenced in a paired-end format on a NovaSeq 6000 (Illumina).

### Bioinformatics Methods

BS and NBS reads were subjected to adaptor trimming (Illumina paired-end sequencing adapters) using Cutadapt v1.2.1 ^52^ with parameter O = 3, and low-quality reads removal using Sickle v1.2 ^53^ with parameters (minimum window quality score > 20 and read length > 15 bp). Quality filtered and adapter trimmed reads were aligned to the NC_009333.1 (NCBI) assembly of the Human herpesvirus 8 strain GK18 genome using BWA-Meth ^54^ with default parameters. RNA modifications (Ψ) were identified on the alignment files using RBSSeqTools ^23^ with the following parameters (Bisulfite reads (BS): ≥ 5 deletions, ≥ 0.02 fraction deletion and ≥ 10 coverage). We merged adjacent positions to form deletion groups, which were pruned to remove positions with less than half the maximum observed fraction deletion in the group. Raw data files available at NCBI GEO (GSE217688).

### Mapping RBS-Seq-identified PSU sites onto KSHV annotated features

KSHV 2.0 annotation ^18^ was merged to NC_009333.1 reference genome through BlastN ^55^ (-FF -W7 - v1 -b1) of all novel features from the former against the latter whole genome. An *ad hoc* PERL script was used to convert Blast tabular format to gff, allowing the integration of both annotations. Gff was converted to bed6 through regular Unix “cut” commands, setting “feature_type” from gff (3^rd^ column) as “feature_name” for bed6 (4^th^ column). Bed files containing BS-Seq-identified PSU sites from the three conditions under investigation (0h, 8h, and 20h) were compared against the recently created NC_009333.1 + KSHV 2.0 bed6 through a “bedtools intersect -loj” execution ^56^.

### PSU motif analysis

STREME tool from the MEME suite ^57^ (--dna --minw 3 --maxw 10 --thresh 0.01) was employed for screening short motifs on the KSHV genome within a 15 nt-long PSU site surrounding area (7 nt upstream - PSU site - 7 nt downstream). PSU sites’ bed files (described above) were combined into a single non-redundant tabular format used as input for an *ad hoc* PERL script aimed at generating the 15 nt-long sequences (with a central PSU site) fasta file used as input for STREME. PSU sites mapped on the minus strand of the genome had their 15 nt-long sequences reversed-complemented. A KSHV whole genome-derived background control file for STREME was created with pyfasta: pyfasta split -n 1 -o 7 -k 15 NC_009333.1.fasta.

### Protein differential enrichment

Proteomics TMT quantification was assessed for differential enrichment with an adapted eBayes function (eb.fit) from the Limma Bioconductor package ^58, 59^ combining both Pus1 and Pus7 experiments against their respective controls. EnhancedVolcano was employed for an overall DE visualisation through volcano plots.

Additionally, our TMT quantification results were compared against a publicly available proteomics dataset (Data are available via ProteomeXchange with identifier PXD037389 ^51^), which made use of SILAC technology for quantifying differentially expressed cellular and KSHV proteins during lytic reactivation. Normalised protein abundance values from both independent experiments were log-scaled and samples were assessed via Spearman Correlation and Hierarchical Clustering analyses. Heat map was plot using the pheatmap R package. All R tools described in this section were run under the R v4.1.0 environment.

### Statistical analysis

Except where otherwise stated, graphical data shown represent mean plus/minus standard error of mean or standard deviation (SD) using at least 3 independent experiments. Differences between means were analysed by Students t-test or one-way ANOVA as described in the figure legends. Statistics were considered significant at p < 0.05 *, p <0.01 **, p <0.001 ***, p <0.0001 **** between groups.

## Data availability

Source data for RBS-Seq and TMT-quantitative mass spectrometry have been deposited to NCBI GEO and PRIDE databases.

## Acknowledgements

We are very thankful to Professor Jae Jung (UCLA) for the TREx BCBL1-Rta cell line and Dr Edwin Chen (University of Westminster) for the psPAX2 and pVSV.G plasmids. We would like to thank Dr. Kate Heesom (Proteomics Facility, University of Bristol, UK) for the proteomics technical service and bioinformatics support. Part of the bioinformatics experiments were undertaken on ARC4, a cluster from the High Performance Computing facility at the University of Leeds, UK. This work was supported in parts by Worldwide Cancer Research (16-1025) and BBSRC (BB/T00021X/1).

## Author Contributions

AW conceived the study and acquired project funding. TJM and KLH performed the experiments and TJM, KLH and AW analysed the resulting data. EJRV and CAA analysed transcriptomics and proteomics datasets. The manuscript draft was written by TJM and AW. All authors reviewed and edited the final version.

## Competing interests

The authors declare no competing financial or non-financial interests.

## Notes

### Competing Interest Statement

The authors have declared no competing interest.

