## Supplementary Figures for "Pseudouridine prevalence in Kaposi’s sarcoma-associated herpesvirus transcriptome reveals an essential mechanism for viral replication"

Supplementary Figures 1-5 and Legends

Supplementary Tables 1-3

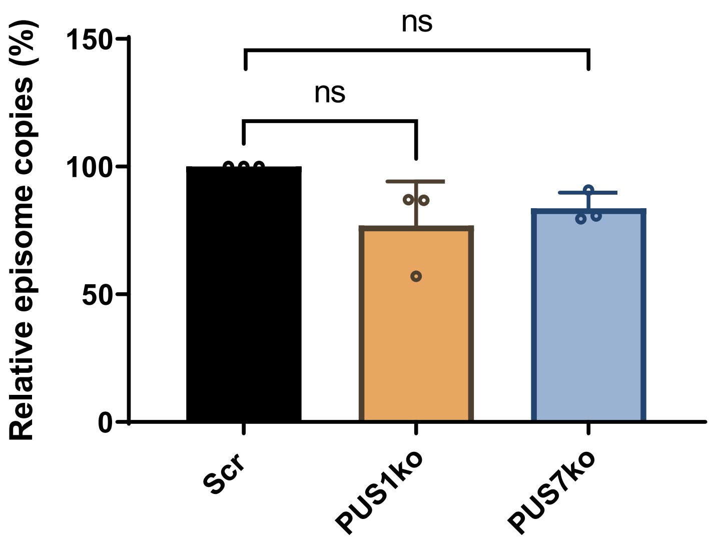


**Supplementary Figure 1. KSHV episome stability in TREx BCBL-Rta PUS1 and PUS7 CRISPR knockout cells.** TREx BCBL1-Rta cells were cultured and samples taken every 7 days. Samples were DNA extracted followed by qRT-PCR. qRT-PCR data was normalized to GAPDH followed by Scrambled control. Error bars represent SD, N = 3 p = <0.05 using a two tailed Students unpaired t-test.

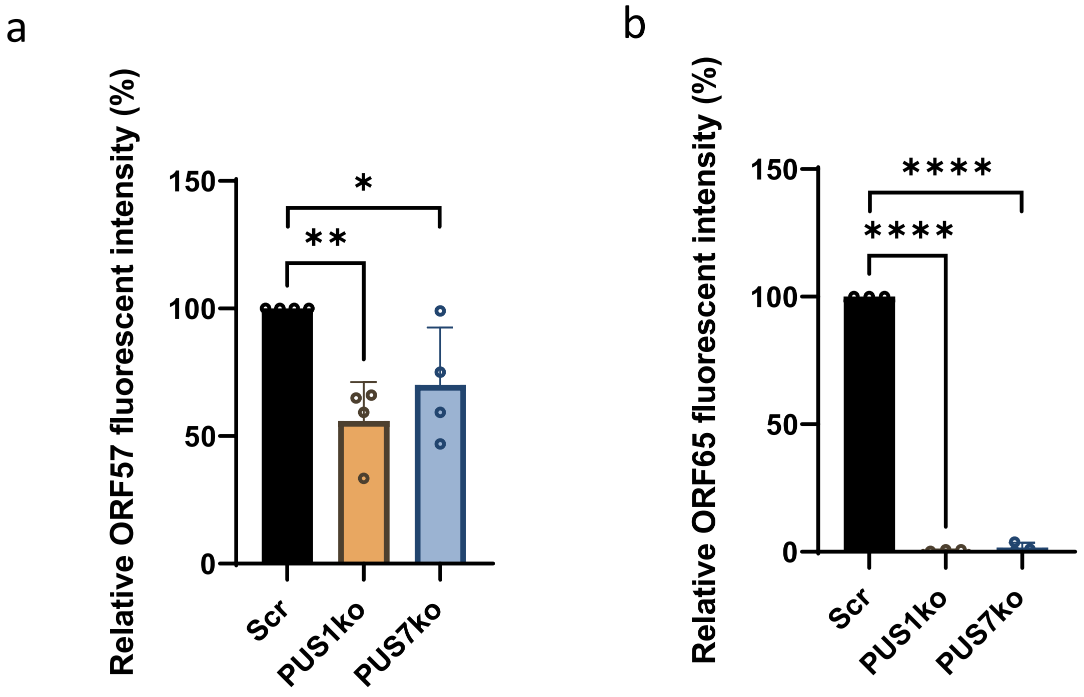


**Supplementary Figure 2. Densitometry of ORF57 and ORF65 TREx BCBL1-RTA PUS1 and PUS7 CRISPR knockout cells during lytic reactivation.** Western blot images from figure 2 were analyzed using Image Studio. Fluorescence data for ORF57 and ORF65 were normalized to Scrambled control. Error bars represent SD, N = 4 p = <0.05 using a two tailed Students unpaired t-test.

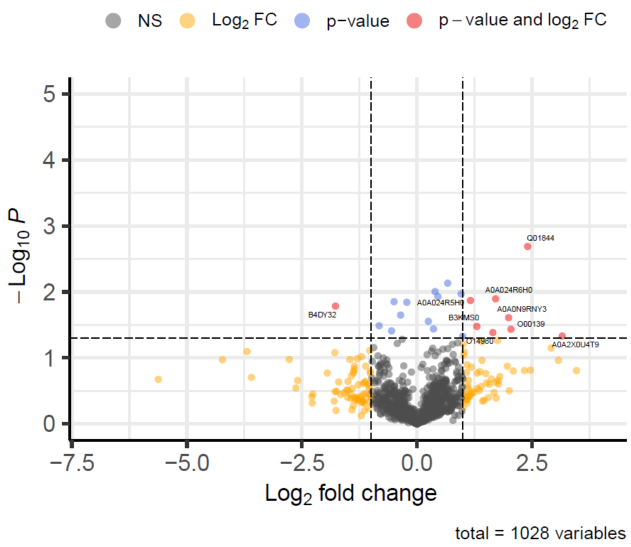


**Supplementary Figure 3. TMT DE analysis volcano plot of PUS1 and PUS7 co-interactors.** Combined PUS1 and PUS7 interactions submitted to eBayes differential enrichment analysis. Each point represents a single protein, plotted by log2 (fold change in abundance) versus –log10(p-value).

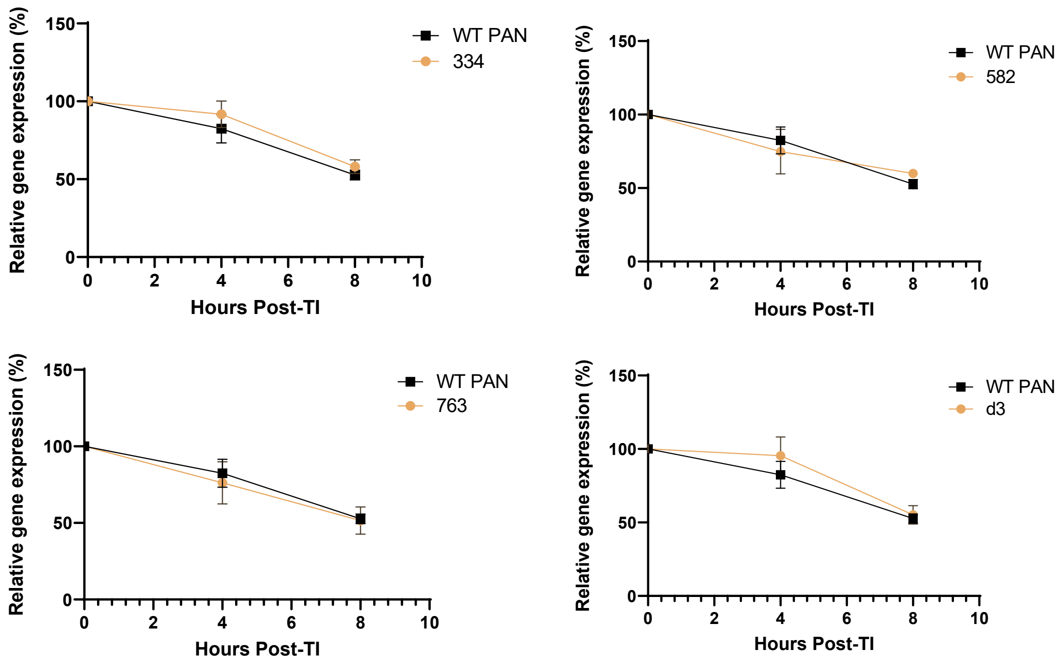


**Supplementary Figure 4. RNA stability of PAN ᵠ mutants without ORF57-eGFP co-transfection.** The stability of PAN mutants 334, 582 and 763 were determined by assessing mRNA decay using actinomycin D (AcD) or control in HEK 293T Cells. Cells were transfected with PAN (or PAN ᵠ mutant) and incubated for 48 hours before the addition of 10 ug/ml AcD or 1% DMSO. Cells were then collected at 2, 4 and 6 hours post AcD treatment and total RNA was extracted followed by qRT-PCR. Error bars represent SE, n = 4 for all experiments

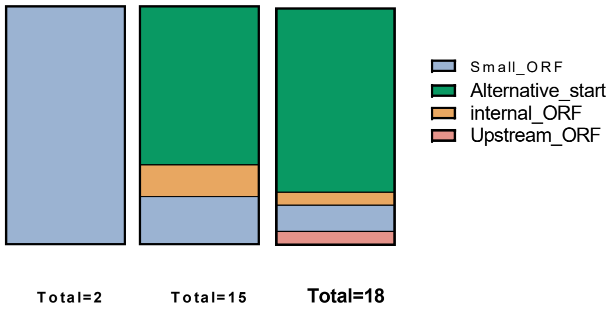


**Supplementary Figure 5. Distribution of ψ sites across alternative topological regions of viral RNA.** Genome features include small ORFs, alternative start ORFs, internal ORFs and upstream ORFs

| Antibody | Usage | Source |
|---|---|---|
| **Anti-ORF57 Mouse** | Western blot, IF | Santa Cruz, sc-135746 |
| **Anti-ORF65 Rabbit** | Western blot | In-house |
| **Anti-PUS1 Rabbit** | Western blot, IF, IP | Sigma, SA131411457 |
| **Anti-PUS7 Mouse** | Western blot, IF, IP | Invitrogen, PA5-54983 |
| **Anti-GAPDH Mouse** | Western blot | Abcam, ab8245 |
| **Anti-Pseudouridine Mouse** | RIP | Diagenode, c15200247 |
| **Anti-IgG Mouse** | IP | Merck, 2430392 |
| **Anti-IgG Rabbit** | IP | Merck, 2679019 |

**Supplementary Table 1. Primary antibodies used in this study**

| Plasmid | Backbone | Source |
|---|---|---|
| **pCMV_PAN** | pCMV | Generated during this study |
| **pCMV_PAN334** | pCMV | Generated during this study |
| **pCMV_PAN582** | pCMV | Generated during this study |
| **pCMV_PAN763** | pCMV | Generated during this study |
| **pCMV_PANΔ3** | pCMV | Generated during this study |
| **lentiCRISPRv2_PUS1_gRNA1** | lentiCRISPRv2. Backbone a gift from Feng Zhang (Addgene plasmid # 52961 ; http://n2t.net/addgene:52961 ; RRID:Addgene_52961) | Generated during this study. |
| **lentiCRISPRv2_PUS7_gRNA1** | lentiCRISPRv2 | Generated during this study |
| **lentiCRISPRv2_Scrambled_gRNA1** | lentiCRISPRv2 | Generated during this study |
| **pCMV_ORF57-eGFP** | pCMV | Previously generated by Whitehouse lab. |

**Supplementary Table 2. Plasmids used and generated in this study.**

| Primer | Sequence 5' – 3' |
|---|---|
| **PAN334_Mut_fwd** | AGGCCAGTTGaAGCCCCCTTT |
| **PAN334_Mut_Rev** | GGAGATTGAATCCAATGCAATAACCC |
| **PAN582_Mut_fwd** | TGGTGCGTTGaGAAGCATTTTAAAATG |
| **PAN582_Mut_Rev** | ATAAGATACACATCCAGATTGTC |
| **PAN763_Mut_fwd** | CTGGGAGCGCaCTTTCAATGTTAATG |
| **PAN763_Mut_Rev** | CTGCCGCACACCACTTTA |
| **PAN_334_CLAP_Forward** | CCCGCAGAACAAAAGCTG |
| **PAN_334_CLAP_Adapter** | pCCATGGCAGCTTTTGTTCTGCGGG |
| **PAN_334_CLAP_Splint** | GTCGCCATGGAGCCCCCTTT/3SpC3/ |
| **PAN_334_CLAP_Reverse** | GCACCACTGTTCTGATACACC |
| **PUS1_gRNA_Fwd** | CACCGGCCCGGACAGACAAGGTGGG |
| **PUS1_gRNA_Rev** | AAACCCCACCTTGTCTGTCCGGGCC |
| **PUS7_gRNA_Fwd** | CACCGGTTGTCGAAGATAATGACAG |
| **PUS7_gRNA_Rev** | AAACCTGTCATTATCTTCGACAACC |
| **PUS1_gRNASeq_Fwd** | GGTGAGTGAGCAGAAAACAG |
| **PUS1_gRNASeq_Rev** | ACAAGAGAAAGCAAAGACCAG |
| **PUS7_gRNASeq_Fwd** | GCTTTTGCCTGGCCGCCCTA |
| **PUS7_gRNASeq_Rev** | CCTCCTCCTCGCACTCCTCTGA |
| **PAN_qPCR_Fwd** | ATAGGCGACAAAGTGAGGTGGCAT |
| **PAN_qPCR_Rev** | TAACATTGAAAGAGCGCTCCCAGC |
| **GAPDH_qPCR_Fwd** | TGTCAGTGGTGGACCTGA |
| **GAPDH_qPCR_Rev** | GTGGTCGTTGAGGGCAATG |
| **ORF57_qPCR_Fwd** | GCCATAATCAAGCGTACTGG |
| **ORF57_qPCR_Rev** | GCAGACAAATATTGCGGTGT |
| **28S_qPCR_Fwd** | CAGGGGAATCCGACTGTTTA |
| **28S_qPCR_Rev** | ATGACGAGGCATTTGGCTAC |
| **ORF4_qPCR_Fwd** | GCCTCAGAGACCGCGAGA |
| **ORF4_qPCR_Rev** | AGCGATTTTTAGACGCCGG |
| **ORF65_qPCR_Fwd** | AAGGTGAGAGACCCCGTGAT |
| **ORF65_qPCR_Rev** | TCCAGGGTATTCATGCGAGC |

**Supplementary Table 3. qPCR and gRNA primers used in this study.**
